# Mpox Virus is Inhibited By Nucleoside Analogues Including the Acyclic Phosphonates Tenofovir and Adefovir

**DOI:** 10.1101/2023.06.30.547277

**Authors:** Jasper Lee, Emerson Ailidh Boggs, Huanchun Zhang, Philip R. Tedbury, Stefan G. Sarafianos

## Abstract

Mpox virus (MPXV) is an orthopoxvirus that causes the human disease mpox, which is characterized by fever, myalgia, and formation of rashes and lesions, and which garnered worldwide attention due to a global outbreak in 2022. In response to the outbreak, the antivirals tecovirimat, cidofovir, and brincidofovir have been used as emergency treatment for mpox. However, because of drug resistance and toxicity risks with those compounds, there is still a need for additional antivirals to treat orthopoxvirus diseases. Since cidofovir is a nucleoside analogue, we investigated a selection of other such compounds for antiviral activity against orthopoxviruses. We developed in vitro screening assays using fluorescent strains of vaccinia virus (VACV) and modified vaccinia Ankara (MVA) to measure the antiviral potency of test compounds. We found that tenofovir alafenamide and adefovir dipixovil, both acyclic phosphonates, had strong potential combinations of anti-orthopoxvirus activity and low toxicity after testing them against MVA and VACV, with EC_50_ values in the single digit micromolar and nanomolar range, while other potential hits included trifluridine and two arabinosides. We then recapitulated the results with MPXV using a luciferase-based assay. These data reinforce the interest of repurposing nucleoside analogues as antivirals to treat poxvirus infections and provide a basis for high throughput screening and mechanistic and antiviral resistance studies.

## INTRODUCTION

Mpox, formerly known as monkeypox, is a disease caused by mpox virus (MPXV) (1). MPXV is classified into the *Orthopoxvirus* genus, which contains double-stranded DNA viruses including as the causative agents of smallpox and cowpox, as well as vaccina virus (VACV). Mpox disease is characterized by fever, myalgia, and lymphadenopathy, followed by the formation of rashes that evolve into lesions and papules all over the body (2). Before 2022, mpox only appeared in sporadic outbreaks in the Democratic Republic of the Congo and Nigeria, where the virus is endemic, although it emerged in the United States in a small outbreak in 2003 (3). Thus, it did not garner much attention until the 2022 global outbreak, which as of April 2023, has reached over 100 countries, with over 85,000 cases worldwide, with the vast majority of which are countries where the virus is not endemic (4). In the United States alone, over 30,000 cases have been confirmed to date. While the number of cases has steadily declined since a peak in early August 2022 (5), this outbreak has demonstrated the potential for future emergence of mpox and other poxviruses and thus the need for vaccines and antivirals to be prepared for the next outbreak, particularly due to the cessation of widespread vaccination efforts against orthopoxviruses following the global eradication of smallpox.

Tecovirimat (TPOXX), cidofovir (CDV), and brincidofovir (BCV) are the three primary antiviral compounds currently available for treating mpox (6). These compounds were selected based on data showing their efficacy in treating smallpox in nonhuman primates (7), with TPOXX and BCV in particular approved by the FDA for this purpose (8, 9). However, treatment data with these compounds for mpox are currently limited. The potential for drug resistance is present in all compounds (10-13). In addition, CDV and BCV have toxicity concerns in humans, in the kidneys and liver respectively (8, 14). Given these factors, there is still a need for discovery and development of new antivirals to treat mpox and other orthopoxviruses, given the low number of effective treatments and the chance for the virus to re-emerge. Here, we tested other nucleoside analogues, particularly acyclic phosphonates, to see if there were any other than CDV that have anti-*Orthopoxvirus* activity, starting with those that have been FDA-approved for treatment against other viruses. We show that out of the compounds tested, adefovir dipixovil, tenofovir alafenamide, and trifluridine have the strongest antiviral potential against VACV and MPXV. In doing so we provide a foundation for further studies with potential *Orthopoxvirus* inhibitors and establish a framework for the discovery of new anti-*Orthopoxvirus* compounds.

## RESULTS

BHK-21, Huh7-C3, and Huh7.5 cells were grown in complete Dulbecco’s modified Eagle’s medium (DMEM) (Corning) supplemented with 10% fetal bovine serum (F2442, Millipore Sigma), 100 U/ml penicillin, 100 μg/ml streptomycin (#400-109, Gemini Bioproducts, West Sacramento, CA, USA), and 2 mM L-glutamine (#25030-081, Gibco), and incubated in a humidified incubator at 37°C with 5% CO_2_.

Tecovirimat, tenofovir alafenamide (TAF), tenofovir disoproxil fumarate (TDF), adefovir, trifluridine, stavudine, and abacavir sulfate were obtained from Selleck Chemicals (Houston, TX, USA). Cidofovir, penciclovir, idoxuridine, Ara-U, Ara-G, were obtained from Cayman Chemical (Ann Abor, MI, USA). Brincidofovir was obtained from AdooQ Bioscience (Irvine, CA, USA). Ara-C, acyclovir, vidarabine (Ara-A), clofarabine, and telbivudine were obtained from Millipore Sigma (Burling, MA, USA). Tenofovir exalidex was obtained from MedKoo Biosciences (Morrisville, NC, USA). Fludarabine was obtained from Medchem Express (Monmouth Junction, NJ, USA). TAF and TDF were obtained from the NIH AIDS reagent program (https://www.hivreagentprogram.org/).

Modified vaccinia Ankara strain VacMVA-p11GFP (MVA-GFP) and vaccinia strain VacWR-p7.5eGFP (VACV-GFP) were obtained from the University of Iowa Viral Vector Core. To screen nucleoside analogues, BHK-21 cells were plated onto 96-well plates, incubated overnight at 37°C with 5% CO_2_, then treated with compounds, before addition of MVA-GFP at an MOI of 0.03. At 48 hours post infection (hpi), plates were analyzed on a Cytation 5 Cell Imaging Multimode Reader (BioTek), where the infected cell count and total fluorescence intensity at were measured using a GFP filter cube with a 469/35 nm excitation filter and a 525/39 nm emission filter, and bright field images were taken using a 4X objective lens to visually observe for possible toxicity (**Figure 1**). Compounds that caused inhibition, shown by a lack of fluorescence compared to positive controls, were then retested in dose response assays, where the raw fluorescence values were used to calculate the EC_50_s of each compound on Prism v.9 (GraphPad), shown in **Table 1**. This assay was first verified using TPOXX, CDV, BCV, and cytarabine (Ara-C), another compound used as a poxvirus DNA synthesis inhibitor (15-17). Then, we selected nucleoside analogues as a major group to test as we had them readily available and the category includes CDV and BCV. Compounds that appeared to cause inhibition and did not appear toxic from visual observations, including both tenofovir formulations, adefovir dipivoxil, Ara-A, Ara-G, trifluridine, and idoxuridine, were selected for further study, firstly through a cytotoxicity check using XTT. For the XTT assay to measure cytotoxicity, 8,000 BHK-21 or 20,000 Huh7-C3 cells were plated on 96-well plates and incubated overnight, then treated with serially diluted compounds. At 48 hpi, XTT was added to plates, which were incubated for 2 h and read at 490/650nm. Raw fluorescence values were used to calculate the CC_50_s of each compound on Prism v.9 (GraphPad) (**Table 1**), showing that Ara-C, TAF, adefovir dipivoxil, and Ara-A were somewhat toxic to BHK-21 cells, but not any of the other possible hits. A number of other compounds were screened and rejected due to lack of inhibition or were clearly toxic to cells at the EC_50_ are found in **Table S1**.

**Table 1.**
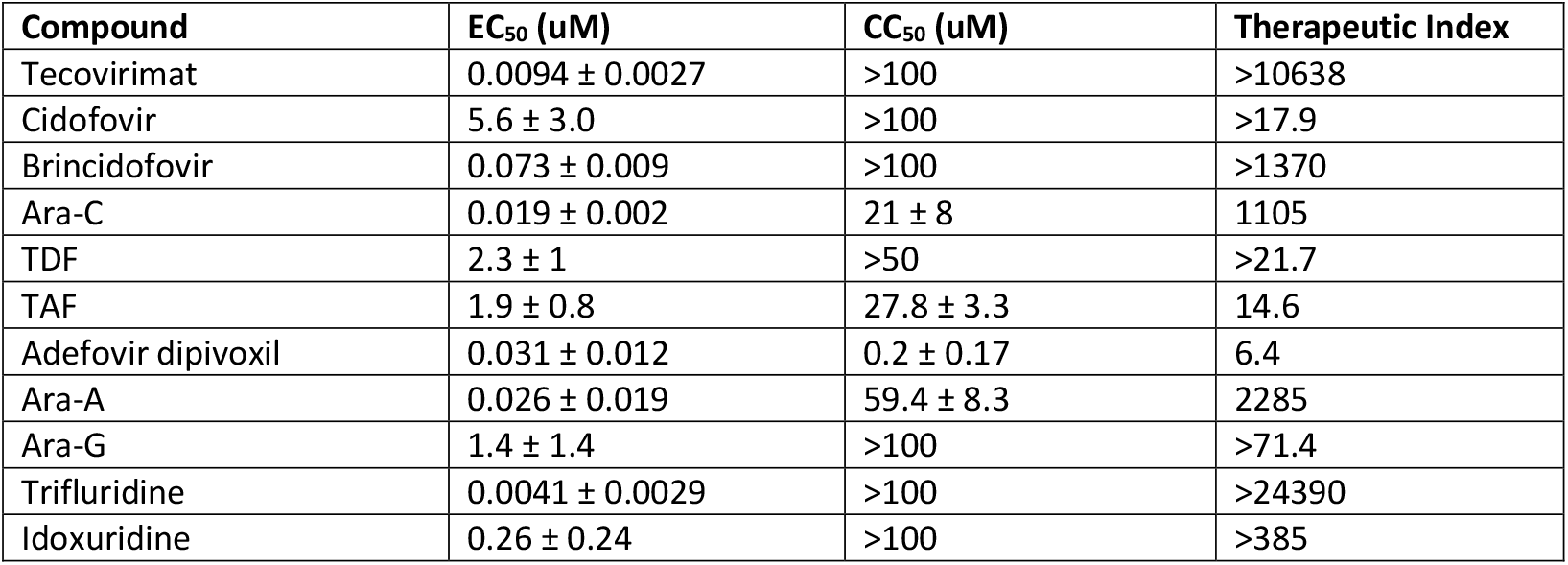
Compounds that showed inhibition of MVA-GFP. BHK-21 cells were seeded in 96-well plates at a concentration of 1 × 10^4^ cells/well and grown overnight, then treated with compound and subsequently infected with MVA-GFP at 0.03 MOI. At 48 hpi, plates were read on a BioTek Cytation 5 where bright field and fluorescence images were taken using the 4X objective. Total fluorescence intensity per well was recorded and used to calculate EC_50_ values, where the EC_50_ was defined as the concentration where there would be a 50% reduction in fluorescence intensity. CC_50_ was determined using XTT to measure cytotoxicity. The therapeutic index was defined as the CC_50_ divided by the EC_50_.

**Figure 1.**
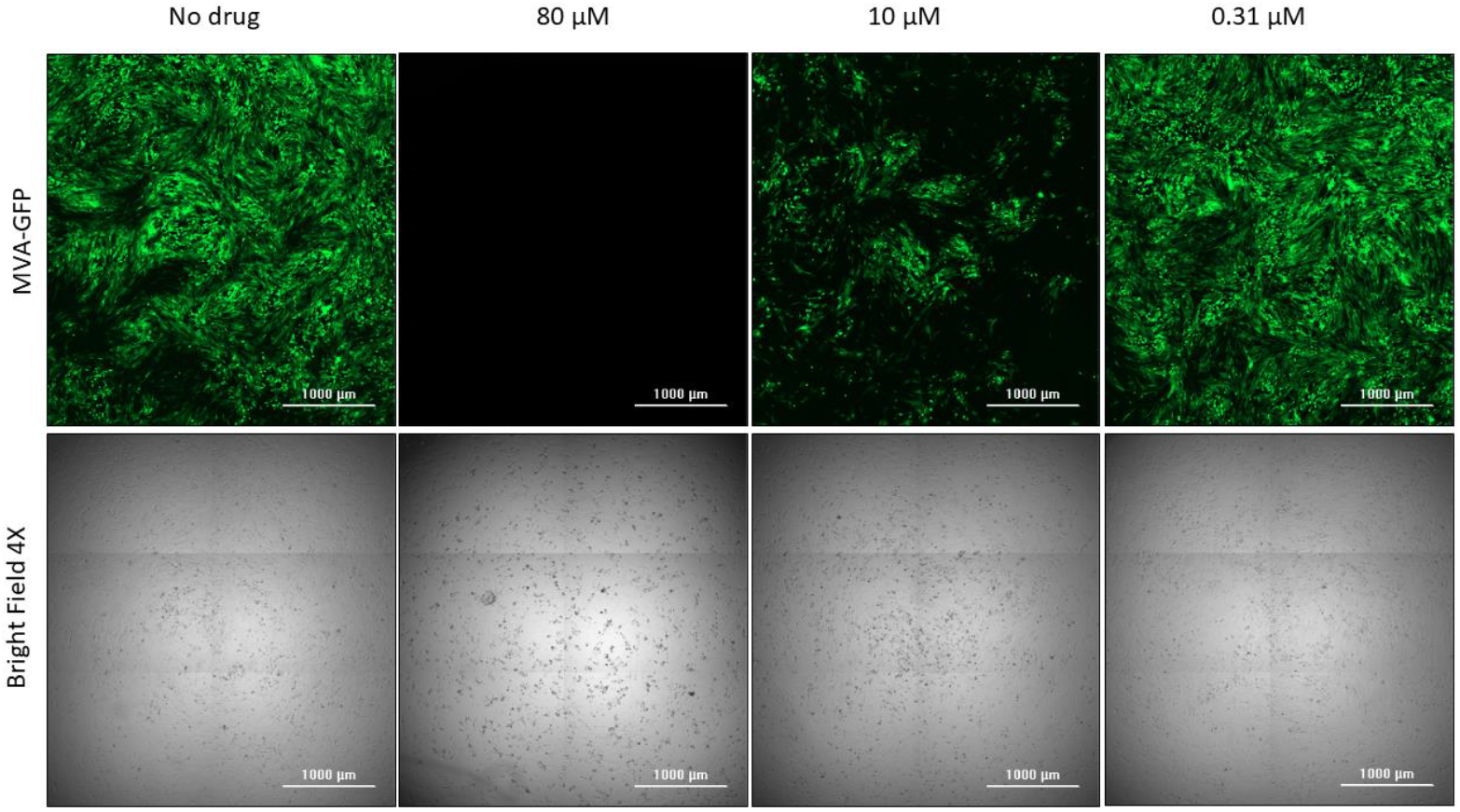
Example fluorescence and bright field images of BHK-21 cells infected with MVA-GFP and treated with TAF in a dose response assay. For each well, 3×2 image montages were taken and stitched together. The top and bottom images are of the same well. Not all tested concentrations are depicted. Scale bar = 1 mm.

Once compounds were identified through screening and dose response assays with MVA-GFP, we re-tested them in dose response assays with VACV-GFP in BHK-21 and Huh7-C3 cells, measuring infected cell count and total fluorescence intensity as with MVA (**Tables 2 and 3, Figure 2**). While there was some variability in the EC_50_ values determined for different viruses using different cell lines, all of them save for TDF were in the low single digit-micromolar or submicromolar range, low enough to continue evaluation. Because we used Huh7-C3 cells, we evaluated cytotoxicity of each compound in that cell line using the XTT assay as described above, showing a lack of toxicity of all compounds in Huh7-C3 cells at the concentrations tested (**Table 3**).

**Table 2.**
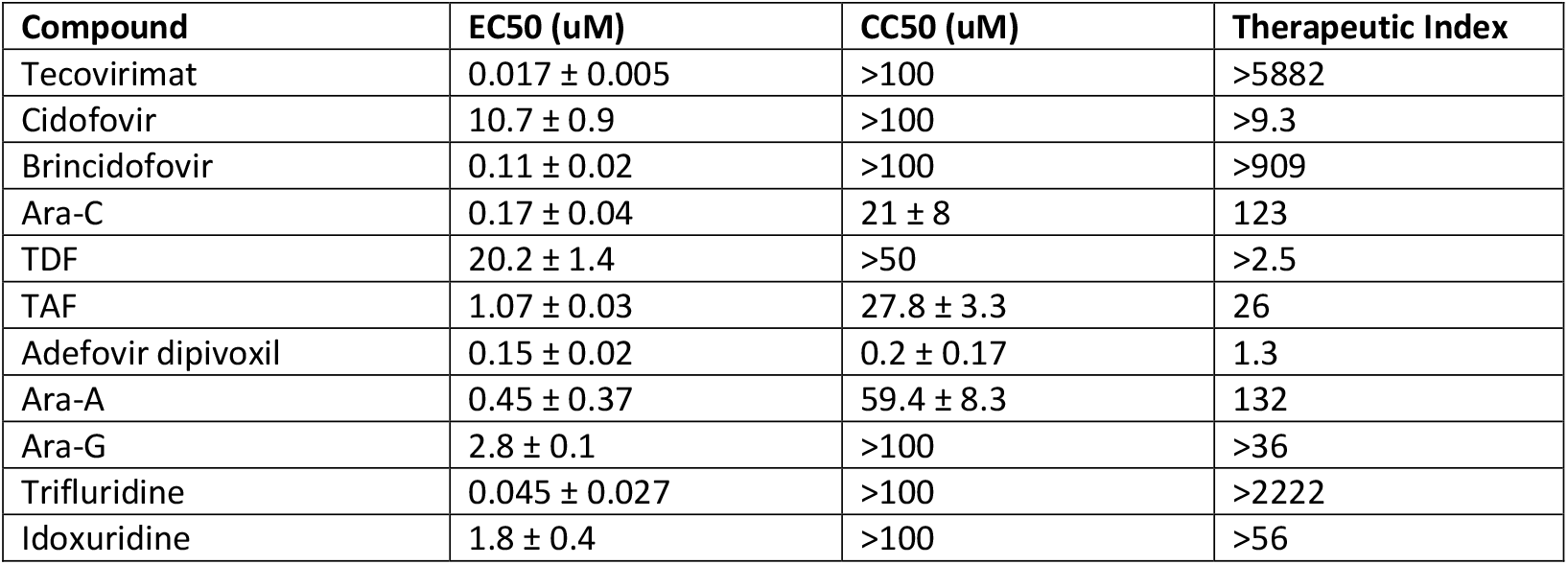
Dose response assays in BHK-21 cells with selected compounds and VACV-GFP. BHK-21 cells were seeded in 96-well plates at a concentration of 1 × 10^4^ cells/well and grown overnight, then treated with compound and subsequently infected with VACV-GFP at 0.06 MOI. At 48 hpi, plates were read on a BioTek Cytation 5 where bright field and fluorescence images were taken using the 4X objective. Total fluorescence intensity per well was recorded and used to calculate EC_50_ values, which were defined as the concentration where there would be a 50% reduction in fluorescence intensity. CC_50_ was determined using XTT to measure cytotoxicity; values here are identical to those listed in Table 1. The therapeutic index was defined as the CC_50_ divided by the EC_50_.

**Table 3.** Dose response assays in Huh7-C3 cells with selected compounds and VACV-GFP. Huh7-C3 cells were seeded in 96-well plates at a concentration of 1 × 10^4^ cells/well and grown overnight, then treated with compound and subsequently infected with VACV-GFP at 0.06 MOI. At 48 hpi, plates were read on a BioTek Cytation 5 where bright field and fluorescence images were taken at 4X. Total fluorescence intensity per well was recorded and used to calculate EC_50_ values, which were defined as the concentration where there would be a 50% reduction in fluorescence intensity. CC_50_ was determined using XTT to measure cytotoxicity. The therapeutic index was defined as the CC_50_ divided by the EC_50_.

| Compound | EC<sub>50</sub> (uM) | CC<sub>50</sub> (uM) | Therapeutic Index |
| --- | --- | --- | --- |
| Tecovirimat | 0.076 ± 0.006 | >100 | >1316 |
| Cidofovir | 6.7 ± 4.2 | >100 | >14.9 |
| Brincidofovir | 0.21 ± 0.13 | >100 | >476 |
| Ara-C | 0.011 ± 0.012 | >50 | >4545 |
| TDF | 29 ± 17 | >100 | >3.4 |
| TAF | 3.7 ± 1.9 | >50 | >13.5 |
| Adefovir dipivoxil | 0.41 ± 0.15 | >50 | >122 |
| Ara-A | 0.20 ± 0.11 | >100 | >500 |
| Ara-G | 0.61 ± 0.72 | >100 | >164 |
| Trifluridine | 0.024 ± 0.002 | >100 | >4167 |
| Idoxuridine | 0.73 ± 0.67 | >100 | >137 |

**Figure 2.**
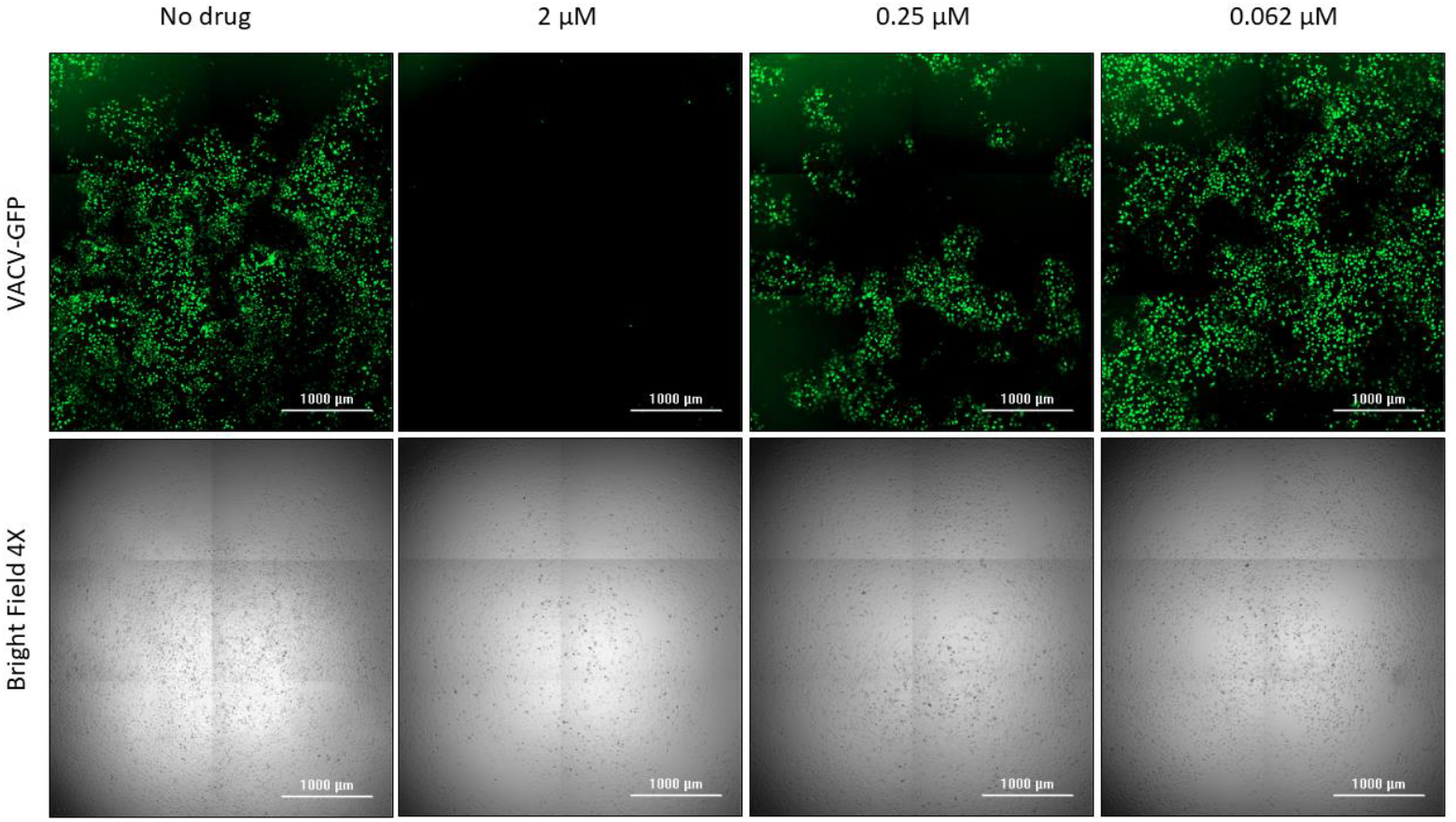
Example fluorescence and bright field images of Huh7-C3 cells infected with VACV-GFP and treated with adefovir dipivoxil in a dose response assay. For each well, 3×2 image montages were taken and stitched together. The top and bottom images are of the same well. Not all tested concentrations are depicted. Scale bar = 1 mm.

Finally, to confirm that tenofovir alafenamide and adefovir dipivoxil were potent against MPXV, we ran dose response assays. MPXV (strain hMPXV/USA/MA001/2022 (Lineage B.1, Clade IIb)) with a firefly luciferase reporter gene (MPXV-Luc) was obtained from Subbian Satheshkumar Panayampalli of the Poxvirus and Rabies Branch in the National Center for Emerging and Zoonotic Infectious Diseases at the Centers for Disease Control and Prevention. MPXV-Luc was used to infect Huh7.5 cells treated with compounds, then incubated for 48 h at 37°C with 5% CO_2_. Afterwards, the plates were analyzed for total luminescence on a BioTek Synergy LX Multi-Mode Microplate Reader, with raw values used to calculate EC_50_ curves and values (**Table 4**). For the tested compounds, we obtained EC_50_ values that were similar to those in the VACV-GFP studies, staying within the range of 1 μM and below.

**Table 4.** Dose response assays in Huh7.5 cells with selected compounds and MPXV-Luc. Huh7.5 cells were seeded in 96-well plates at a concentration of 1 × 10^4^ cells/well and grown overnight, then treated with compound and subsequently infected with MPXV-Luc. At 48 hpi, plates were read on a BioTek Synergy LX Multi-Mode Microplate Reader measuring total luminescence per well, values of which used to calculate EC_50_ values, which were defined as the concentration where there would be a 50% reduction in luminescence intensity.

| Compound | EC<sub>50</sub> (uM) |
| --- | --- |
| Tecovirimat | 0.45 ± 0.74 |
| Cidofovir | 8.6 ± 0.5 |
| Ara-C | 0.34 ± 0.39 |
| TAF | 1.2 ± 0.2 |
| Adefovir dipivoxil | 0.35 ± 0.05 |
| Ara-A | 0.81 ± 0.53 |
| Ara-G | 1.9 ± 0.1 |

## DISCUSSION

In this study, we aimed to establish a streamlined protocol for screening and identifying compounds that inhibit MPXV, starting with nucleoside analogues. Our data show the potential for nucleoside analogues, particularly acyclic phosphonates, in the inhibition of orthopoxvirus spread. In addition to determining EC_50_ values for tecovirimat and cidofovir that align with previous studies (18-20), we also show other potential candidates for further examination. These compounds include adefovir dipivoxil and tenofovir alafenamide, prodrugs of adefovir and tenofovir respectively. Adefovir is used to treat hepatitis B virus (HBV) infection (21), while tenofovir is used to treat both HBV and HIV-1 (22, 23). The efficacy of tenofovir is of particular interest as during the 2022 mpox outbreak, a significant number of cases were found in patients with HIV-1, which raises the possibility that HIV-1 patients currently treated with TAF may benefit from a protective effect of TAF against MPXV. For both compounds, however, further study is obviously needed as the emergence of resistance is a potential concern, as has been shown when used to treat chronic hepatitis B infections (24) and through *in vitro* studies (25). Another inhibitor, trifluridine, was also found to have efficacy. It is primarily used to treat herpesvirus infections, particularly in the eye, but has seen some use in treating vaccinia keratitis in the eye, and in one case, as part of an mpox treatment; therefore, its efficacy in our experiments is consistent with previous relevant reports (26, 27).

This appears to be the first instance in which tenofovir has been shown to inhibit Mpox virus in a cell culture system. Most recently, a comparable potency was reported in a related system for inhibition of VACV by adefovir dipivoxil (28), providing additional evidence that it merits further testing in animals and for the potential for antiviral resistance to develop, which to our knowledge have not been studied. In addition, the potential advantage of tenofovir and adefovir is that they are already being taken by individuals that are at high risk of contracting mpox. In conclusion, we believe this study provides a relatively rapid, concise test for future antiviral discovery that can be further improved by future miniaturization of the assay towards an ultra-high throughput screening approach to identify additional compounds for treating mpox and be a springboard for resistance and mechanistic studies.

## ACKNOWLEDGEMENTS

We would like to thank the Viral Vector Core at the University of Iowa and Subbian Satheshkumar Panayampalli of the Poxvirus and Rabies Branch in the National Center for Emerging and Zoonotic Infectious Diseases at the Centers for Disease Control and Prevention for providing us the viruses used in these experiments. We also thank Dan Loeb, U. Wisconsin for the Huh7-C3 cells and Raymond F. Schinazi and his group for useful interactions in the early stages of the project. SGS acknowledges funding from the Nahmias-Schinazi Distinguished Chair in Research and from the Imagine, Innovate, and Impact Award, I3RAPIDSynergy2022, from Emory School of Medicine and Georgia Clinical & Translational Science Alliance.

---

**Table S1.**
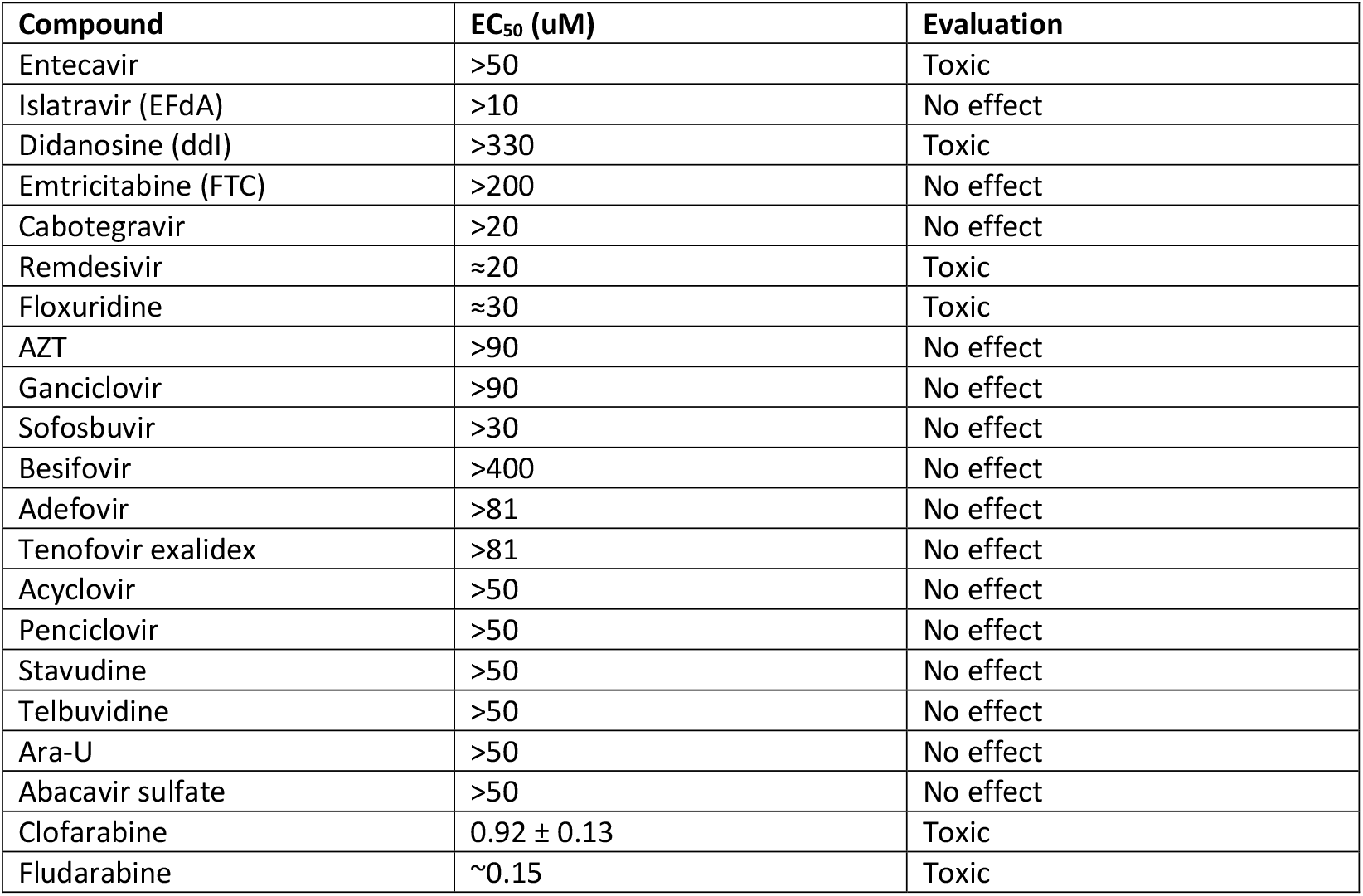
Compounds that were rejected after screening due to lack of inhibition or visible toxicity towards cells at the calculated or estimated EC_50_. BHK-21 cells were seeded in 96-well plates at a concentration of 1 × 10^4^ cells/well and grown overnight, then treated with compound and subsequently infected with MVA-GFP at 0.03 MOI. At 48 hpi, plates were read on a BioTek Cytation 5 where bright field and fluorescence images were taken at 4X. Total fluorescence intensity per well was recorded and used to calculate EC_50_ values where possible. The EC_50_ was defined as the concentration where there would be a 50% reduction in fluorescence intensity.

| Compound | EC<sub>50</sub> (uM) | Evaluation |
| --- | --- | --- |
| Entecavir | >50 | Toxic |
| Islatravir (EFdA) | >10 | No effect |
| Didanosine (ddI) | >330 | Toxic |
| Emtricitabine (FTC) | >200 | No effect |
| Cabotegravir | >20 | No effect |
| Remdesivir | ≈20 | Toxic |
| Floxuridine | ≈30 | Toxic |
| AZT | >90 | No effect |
| Ganciclovir | >90 | No effect |
| Sofosbuvir | >30 | No effect |
| Besifovir | >400 | No effect |
| Adefovir | >81 | No effect |
| Tenofovir exalidex | >81 | No effect |
| Acyclovir | >50 | No effect |
| Penciclovir | >50 | No effect |
| Stavudine | >50 | No effect |
| Telbuvudine | >50 | No effect |
| Ara-U | >50 | No effect |
| Abacavir sulfate | >50 | No effect |
| Clofarabine | 0.92 ± 0.13 | Toxic |
| Fludarabine | ~0.15 | Toxic |

## REFERENCES

1. World Health Organization. 2022. WHO recommends new name for monkeypox disease, on World Health Organization. https://www.who.int/news/item/28-11-2022-who-recommends-new-name-for-monkeypox-disease. Accessed January 19.

2. Philpott D, Hughes CM, Alroy KA, Kerins JL, Pavlick J, Asbel L, Crawley A, Newman AP, Spencer H, Feldpausch A, Cogswell K, Davis KR, Chen J, Henderson T, Murphy K, Barnes M, Hopkins B, Fill M-MA, Mangla AT, Perella D, Barnes A, Hughes S, Griffith J, Berns AL, Milroy L, Blake H, Sievers MM, Marzan-Rodriguez M, Tori M, Black SR, Kopping E, Ruberto I, Maxted A, Sharma A, Tarter K, Jones SA, White B, Chatelain R, Russo M, Gillani S, Bornstein E, White SL, Johnson SA, Ortega E, Saathoff-Huber L, Syed A, Wills A, Anderson BJ, Oster AM, Christie A, et al. 2022. Epidemiologic and Clinical Characteristics of Monkeypox Cases - United States, May 17-July 22, 2022. MMWR Morbidity and mortality weekly report. 71(32):1018–1022. doi:10.15585/mmwr.mm7132e3.

3. US Centers for Disease Control and Prevention. 2003. Multistate outbreak of monkeypox--Illinois, Indiana, and Wisconsin, 2003. MMWR Morb Mortal Wkly Rep 52:537–40.

4. Centers for Disease Control and Prevention (CDC). 2022. 2022 Monkeypox Outbreak Global Map. https://www.cdc.gov/poxvirus/mpox/response/2022/world-map.html. Accessed April 7.

5. Centers for Disease Control and Prevention (CDC). 2022. U.S. Mpox Case Trends Reported to CDC. https://www.cdc.gov/poxvirus/mpox/response/2022/mpx-trends.html. Accessed April 7.

6. Matias WR, Koshy JM, Nagami EH, Kovac V, Moeng LR, Shenoy ES, Hooper DC, Madoff LC, Barshak MB, Johnson JA, Rowley CF, Julg B, Hohmann EL, Lazarus JE. 2022. Tecovirimat for the Treatment of Human Monkeypox: An Initial Series From Massachusetts, United States. Open Forum Infectious Diseases 9.

7. Huggins J, Goff A, Hensley L, Mucker E, Shamblin J, Wlazlowski C, Johnson W, Chapman J, Larsen T, Twenhafel N, Karem K, Damon IK, Byrd CM, Bolken TC, Jordan R, Hruby D. 2009. Nonhuman primates are protected from smallpox virus or monkeypox virus challenges by the antiviral drug ST-246. Antimicrob Agents Chemother 53:2620–5.

8. US Food and Drug Administration. 2021. FDA approves drug to treat smallpox. https://www.fda.gov/drugs/news-events-human-drugs/fda-approves-drug-treat-smallpox. Accessed January 17.

9. US Food and Drug Administration. 2018. FDA approves the first drug with an indication for treatment of smallpox. https://www.fda.gov/news-events/press-announcements/fda-approves-first-drug-indication-treatment-smallpox. Accessed January 17.

10. U.S. Food & Drug Administration (FDA). 2022. FDA Monkeypox Response. https://www.fda.gov/emergency-preparedness-and-response/mcm-issues/fda-mpox-response. Accessed 03/07/2023.

11. Duraffour S, Lorenzo MM, Zöller G, Topalis D, Grosenbach D, Hruby DE, Andrei G, Blasco R, Meyer H, Snoeck R. 2015. ST-246 is a key antiviral to inhibit the viral F13L phospholipase, one of the essential proteins for orthopoxvirus wrapping. Journal of Antimicrobial Chemotherapy 70:1367–1380.

12. Alarcón J, Kim M, Terashita D, Davar K, Garrigues JM, Guccione JP, Evans MG, Hemarajata P, Wald-Dickler N, Holtom P, Garcia Tome R, Anyanwu L, Shah NK, Miller M, Smith T, Matheny A, Davidson W, Hutson CL, Lucas J, Ukpo OC, Green NM, Balter SE. 2023. An Mpox-Related Death in the United States. New England Journal of Medicine 388:1246–1247.

13. Andrei G, Gammon DB, Fiten P, De Clercq E, Opdenakker G, Snoeck R, Evans DH. 2006. Cidofovir resistance in vaccinia virus is linked to diminished virulence in mice. J Virol 80:9391–401.

14. Adler H, Gould S, Hine P, Snell LB, Wong W, Houlihan CF, Osborne JC, Rampling T, Beadsworth MBJ, Duncan CJA, Dunning J, Fletcher TE, Hunter ER, Jacobs M, Khoo SH, Newsholme W, Porter D, Porter RJ, Ratcliffe L, Schmid ML, Semple MG, Tunbridge AJ, Wingfield T, Price NM, Abouyannis M, Al-Balushi A, Aston S, Ball R, Beeching NJ, Blanchard TJ, Carlin F, Davies G, Gillespie A, Hicks SR, Hoyle M-C, Ilozue C, Mair L, Marshall S, Neary A, Nsutebu E, Parker S, Ryan H, Turtle L, Smith C, van Aartsen J, Walker NF, Woolley S, Chawla A, Hart I, Smielewska A, et al. 2022. Clinical features and management of human monkeypox: a retrospective observational study in the UK. The Lancet Infectious Diseases 22:1153–1162.

15. Weir JP, Moss B. 1987. Determination of the transcriptional regulatory region of a vaccinia virus late gene. Journal of Virology 61:75–80.

16. Myskiw C, Piper J, Huzarewich R, Booth TF, Cao J, He R. 2010. Nigericin is a potent inhibitor of the early stage of vaccinia virus replication. Antiviral Research 88:304–310.

17. Yang Z, Maruri-Avidal L, Sisler J, Stuart CA, Moss B. 2013. Cascade regulation of vaccinia virus gene expression is modulated by multistage promoters. Virology 447:213–220.

18. Frenois-Veyrat G, Gallardo F, Gorge O, Marcheteau E, Ferraris O, Baidaliuk A, Favier AL, Enfroy C, Holy X, Lourenco J, Khoury R, Nolent F, Grosenbach DW, Hruby DE, Ferrier A, Iseni F, Simon-Loriere E, Tournier JN. 2022. Tecovirimat is effective against human monkeypox virus in vitro at nanomolar concentrations. Nat Microbiol 7:1951–1955.

19. Yang G, Pevear DC, Davies MH, Collett MS, Bailey T, Rippen S, Barone L, Burns C, Rhodes G, Tohan S, Huggins JW, Baker RO, Buller RL, Touchette E, Waller K, Schriewer J, Neyts J, DeClercq E, Jones K, Hruby D, Jordan R. 2005. An orally bioavailable antipoxvirus compound (ST-246) inhibits extracellular virus formation and protects mice from lethal orthopoxvirus Challenge. J Virol 79:13139–49.

20. Kornbluth RS, Smee DF, Sidwell RW, Snarsky V, Evans DH, Hostetler KY. 2006. Mutations in the E9L polymerase gene of cidofovir-resistant vaccinia virus strain WR are associated with the drug resistance phenotype. Antimicrob Agents Chemother 50:4038–43.

21. Yuen M-F, Lai C-L. 2004. Adefovir dipivoxil in chronic hepatitis B infection. Expert Opinion on Pharmacotherapy 5:2361–2367.

22. Abdul Basit S, Dawood A, Ryan J, Gish R. 2017. Tenofovir alafenamide for the treatment of chronic hepatitis B virus infection. Expert Review of Clinical Pharmacology 10:707–716.

23. Ray AS, Fordyce MW, Hitchcock MJM. 2016. Tenofovir alafenamide: A novel prodrug of tenofovir for the treatment of Human Immunodeficiency Virus. Antiviral Research 125:63–70.

24. Lee Y-S, Suh DJ, Lim Y-S, Jung SW, Kim KM, Lee HC, Chung Y-H, Lee YS, Yoo W, Kim S-O. 2006. Increased risk of adefovir resistance in patients with lamivudine-resistant chronic hepatitis B after 48 weeks of adefovir dipivoxil monotherapy. Hepatology 43:1385–1391.

25. Margot Nicolas A, Johnson A, Miller Michael D, Callebaut C. 2015. Characterization of HIV-1 Resistance to Tenofovir Alafenamide In Vitro. Antimicrobial Agents and Chemotherapy 59:5917–5924.

26. Altmann S, Brandt CR, Murphy CJ, Patnaikuni R, Takla T, Toomey M, Nesbit B, McIntyre K, Covert J, Dubielzig R, Leatherberry G, Adkins E, Kodihalli S. 2011. Evaluation of Therapeutic Interventions for Vaccinia Virus Keratitis. The Journal of Infectious Diseases 203:683–690.

27. Perzia B, Theotoka D, Li K, Moss E, Matesva M, Gill M, Kibe M, Chow J, Green S. 2023. Treatment of ocular-involving monkeypox virus with topical trifluridine and oral tecovirimat in the 2022 monkeypox virus outbreak. American Journal of Ophthalmology Case Reports 29:101779.

28. Dsouza L, Pant A, Offei S, Priyamvada L, Pope B, Satheshkumar PS, Wang Z, Yang Z. 2023. Antiviral activities of two nucleos(t)ide analogs against vaccinia, mpox, and cowpox viruses in primary human fibroblasts. Antiviral Research 216:105651.

